# A bioorthogonal antibody-based chemically-induced-dimerization switch for therapeutic application

**DOI:** 10.1101/2023.04.11.536272

**Authors:** Alexander J. Martinko, Hai L. Tran, Erin F. Simonds, Allison L. Cooke, Zhong Huang, Sruthi Raguveer, Judit Pina Agullet, Suchitra Prasad, Catherine Going, Lisa Marshall, Timothy Park, Sunandan Banerjee, Ramsay Macdonald, Mike Jian, Kenneth Ng, Akhila Palakodaty, Manpreet Kaur, Alberto Ponce, Mohammad Tabrizi, Zachary B. Hill

**Affiliations:** Soteria Biotherapeutics, San Mateo, California, USA

**Author notes:** Both Authors Contributed Equally to this Work (Alexander J. Martinko, Hai L. Tran).

## Abstract

We present the Indinavir Ligand Induced Transient Engagement switch (IDV LITE Switch), a fully synthetic Chemically Induced Dimerization (CID) system wherein two humanized antibody fragments are heterodimerized by the antiviral drug indinavir. The IDV LITE Switch represents the first CID system made from fully humanized protein components and dimerized by a clinically approved small molecule drug lacking a mammalian target, making it an ideal bioorthogonal molecular switch for application in small-molecule controlled therapeutics.

---

Chemically induced dimerization (CID) is an established approach that utilizes a small molecule to regulate the proximity of proteins of interest with precise temporal control. Since their original development in the early 1990s, CID systems have been powerful research tools to manipulate and interrogate signaling events and transcriptional programs dependent on proximity ^1 2^. In addition to the numerous uses of CIDs as research tools, application of CIDs to therapeutics for small molecule regulated activity, including on-off and safety switches, has been explored across gene therapies, cell therapies, and antibody therapeutics ^3^. Notable examples include Ariad Pharmaceuticals’ ARGENT system for small-molecule-controlled gene expression, Bellicum Pharmaceuticals’ inducible caspase-9 safety switch for stem cell transplantation, and Cell Design Labs’ implementation of CIDs to control CAR-T cell therapies ^4 5 6^. Despite nearly 30 years of development of CIDs, currently available tools are mainly derived from naturally occurring systems and often lack bioorthogonality, utilize small molecules with poor drug-like properties, or are commonly built from non-human proteins with high immunogenicity risk. Collectively, these limitations limit the broader utility of CID technology in medicine.

Recently, Foight & Wang described a novel CID system utilizing clinically approved HCV-protease inhibitors ^7^. While this technology uses a bioorthogonal small molecule with drug like properties, the viral-derived nature of one protein domain, and the *de novo* designed nature of the other may lead to immunogenicity issues if incorporated into human therapies. We recently described a method to generate novel CID systems, termed Ligand Induced Transient Engagement Switches (LITE Switches), by raising chemical-epitope-selective antibodies that recognize small molecule drugs in complex with their human protein drug targets ^8 9^. However, this approach is limited in that the small molecules inherently lack biorthogonality due to the pharmacology linked to binding of the intended human protein target. We postulated that we could potentially extend this approach to antiviral small molecules but also maintain exclusively human protein components by engineering both protein components from antibodies. Here we describe the Indinavir LITE Switch, a synthetic CID system built entirely from human antibody fragments and dimerized by the clinically approved antiviral drug, indinavir. To our knowledge, this molecular switch is the first CID described to date that utilizes fully humanized protein components in combination with a clinically approved small molecule drug lacking a mammalian target. The IDV LITE Switch is bioorthogonal, demonstrates practical on/off kinetics *in vivo*, and presents low risk of immunogenicity. Thus, we believe the IDV LITE Switch described here represents an ideal CID for application to human therapeutics.

We hypothesized that the first-generation HIV protease inhibitor, indinavir, could serve as an optimal dimerization agent for a therapeutically applicable CID. Indinavir is an FDA-approved drug that has been given to thousands of patients at high doses, often for periods of years, and is generally well tolerated due to its lack of an intended human protein target ^10^. Importantly, indinavir is metabolized on the order of hours (elimination half-life: 1.8 ± 0.4 hrs) which could facilitate the ability to rapidly and reversibly control CID assembly *in vivo* ^11^. Finally, indinavir is off-patent and synthetic routes to generate low-cost research-grade as well as cGMP indinavir are readily available ^12^.

When considering protein scaffolds as an engineering starting point, we ultimately turned to the most therapeutically experienced proteins known, antibodies. Antibodies are readily engineered, amenable to manufacturing at a therapeutic scale, and can be humanized to mitigate risk of immunogenicity ^13^. Moreover, antibody fragments are highly modular and the ability to fuse antibody fragments to proteins of interest in various orientations and geometries both intra- and extra-cellularly is well established ^14^.

In order to construct the IDV LITE Switch, we sequentially engineered each antibody fragment (Fig.1a). To generate the first antibody fragment for the IDV LITE Switch (LSA), mice were immunized with indinavir conjugated to the carrier protein Keyhole Lymphocyte Hemocyanin (KLH) (Supplementary Fig.S1). An immune phage library was constructed from splenocytes of immunized mice and panned for binders to indinavir conjugated to bovine serum albumin (BSA). Hits were then screened for binding to biotin-indinavir by bio-layer interferometry (BLI) and competitive inhibition of that binding with soluble indinavir (Supplementary Fig.S2). Ab15 was selected for development, was humanized, and underwent affinity and stability engineering to result in the antibody fragment termed “LSA”, which had a measured affinity of 4.6 nM to free indinavir (Fig.1b).

**Fig. 1.**
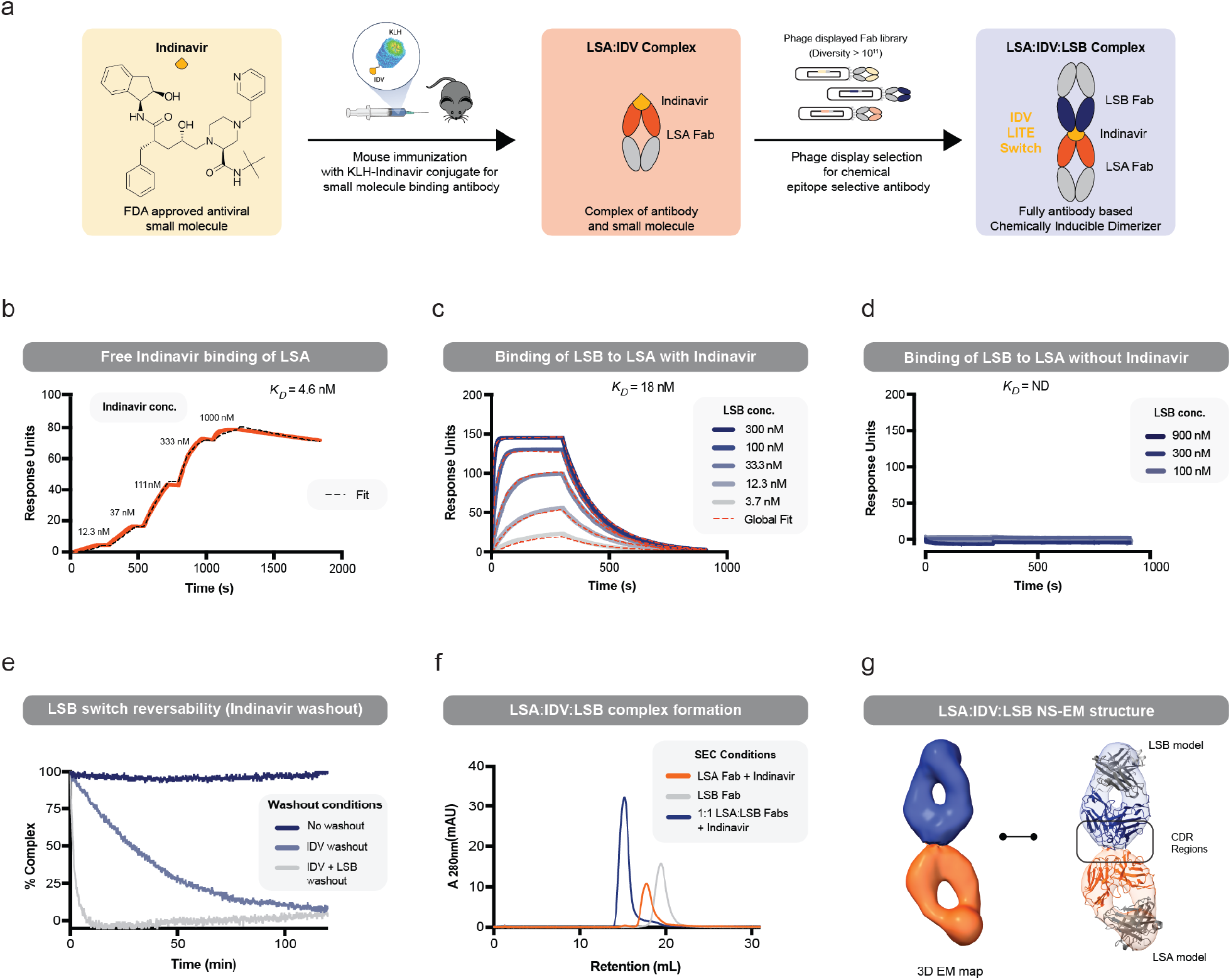
Development of a human antibody-based Indinavir LITE switch. **a**, An antibody discovery campaign against indinavir-KLH followed by humanization led to the development of the indinavir-binding humanized antibody fragment, LSA. A synthetic human Fab-phage library was used to discover a binder specific to the LSA:indinavir chemical epitope, LSB. **b**, SPR single-cycle kinetics of free indinavir binding to immobilized LSA. **c and d**, SPR multi-cycle kinetics of LSB binding to immobilized LSA in the presence and absence of 10µM indinavir. **e**, BLI complex reversibility experiment where the biotin-LSA:indinavir:LSB complex was formed on a streptavidin biosensor and indinavir or indinavir+LSB was washed out, demonstrating near complete reversibility within 2h or 10min, respectively. **f**, SEC chromatogram LSA Fab + indinavir, LSB Fab, and a 1:1 mixture LSA:LSB Fabs + indinavir showing complex formation in solution. **g**, A 3D EM map of the LSA:indinavir:LSB complex generated from 50K particles from negative stain EM. Antibody models of LSA (orange) and LSB (blue) were fit into the density showing CDR regions of both antibodies making contact.

To construct the second half of the IDV LITE Switch, we set out to discover Fab fragments that selectively bound to the LSA:indinavir complex. We utilized a phage displayed human Fab library and applied our previously described phage selection approach to eliminate binders to LSA in the absence of indinavir and enrich for binders that conditionally bind LSA in the presence of indinavir ^8^. One such binder was identified and subsequently engineered to reduce hydrophobicity (to improve developability) and weaken affinity (to improve complex reversibility), yielding the antibody fragment termed “LSB”. LSB had a measured affinity of 18 nM to LSA in the presence of excess indinavir (Fig.1c) and no observable binding in the absence of indinavir (Fig.1d). This data demonstrates indinavir-dependent formation of the LSA/LSB LITE Switch complex.

The IDV LITE Switch is a non-covalent complex, and thus would be expected to disassociate on a timeframe of minutes to hours when indinavir is removed. A BLI assay was performed to assess the reversibility kinetics of the LS2A/LS2B complex upon indinavir washout (Fig.1e). Biotinylated-LSA was immobilized on streptavidin sensors and the LITE Switch complex was formed by adding 1 µM indinavir and 50 nM LSB. Data showed that when concentrations of LSB and indinavir remained constant, no LITE Switch dissociation was observed. In contrast, within 2 hours of indinavir washout, the LITE Switch dissociated to nearly 0% complex, supporting the reversible formation of the heterodimer.

We hypothesized that the LITE Switch assembled as a 1:1 heterodimer in the presence in indinavir. To test this, purified LSA and LSB Fab fragments were assessed by analytical size exclusion chromatography (SEC) alone and in a 1:1 mixture with saturating indinavir (Fig.1f). The earlier retention time of the mixture and absence of monomer peaks suggests that a higher molecular weight dimeric complex was formed. To further confirm the formation of a 1:1 complex, we applied negative stain electron microscopy to determine the structure of the LITE Switch complex formed by LSA, LSB, and indinavir (Supplementary Fig.S4). Images of 54.6K particles were used to determine 2D class averages with 92% representing Fab-Fab-indinavir complexes and 8% representing individual Fabs. A 14.6Å 3D map was calculated from 50K particles and models of LSA and LSB Fabs were fit into the density demonstrating that the LITE Switch complex forms a 1:1 heterodimeric complex wherein the CDR regions of LSA and LSB come into direct contact (Fig.1g). This data, taken together with the other biophysical characterization, collectively supports that the system behaves as a reversibly formed indinavir-inducible antibody-based heterodimer, mechanistically consistent with the intended design.

To demonstrate proof of concept of the IDV LITE Switch to tunably control the activity of a therapeutic modality, we applied the CID technology to regulate the activity of a T-LITE, a small-molecule inducible bispecific T-cell engager format that we have described previously (Fig.2a) ^9^. Here, we segregated the αCD3 (αT-cell) and αEpCAM (αTumor Associated Antigen) arms of a T-cell engager into two inactive bispecifics each incorporating half of the IDV LITE Switch such that they can be dimerized and functionally activated to drive T-cell mediated cell killing with the addition of indinavir.

**Fig. 2.**
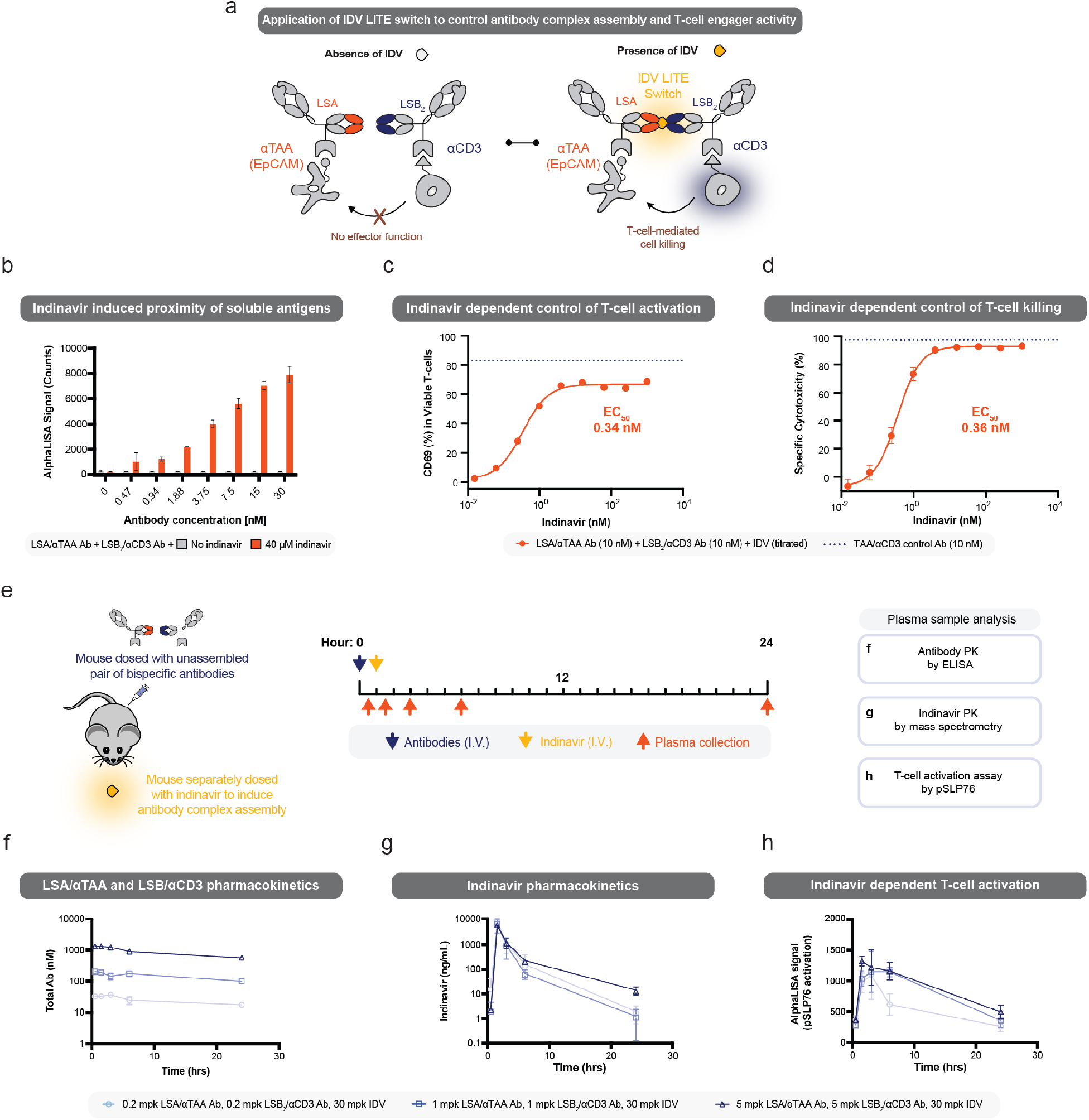
Application of the LITE switch to control the assembly and activity of a T-cell engager. **A**, Schematic representation of the T-LITE platform where indinavir can assemble a functional T-cell engager by bringing together LSA/αTAA and LSB/αCD3 bispecific antibodies. **b**, An AlphaLISA assay demonstrating that the T-LITE platform can bring soluble CD3 and TAA proteins in proximity in an indinavir dose-dependent manner. **c and d**, A TDCC co-culture assay demonstrating the indinavir dose-dependent activation of T-cells and T-cell mediated killing of target cells. **e, f, g, and h**, Study design and pharmacokinetics of T-LITE pairs at three doses with indinavir demonstrating slow clearance of the antibodies and rapid clearance of indinavir in 24hr in C57BL/6 mice. Plasma samples were taken and analyzed in an *ex vivo* pSLP76 assay to measure the level of T-cell activation. Data presented here shows a correlation between T-cell activation and indinavir PK.

A cell-free AlphaLISA assay was developed to quantify the amount of soluble CD3 and EpCAM proteins brought together in close proximity (Supplementary Fig.S5). This assay was used to demonstrate that the indinavir LITE Switch could drive the proximity of CD3 and EpCAM together in solution in an indinavir dose-dependent manner (Fig.2b). Next, we examined assembly of a functional T-LITE T-Cell Engager in a T-cell dependent cellular cytotoxicity (TDCC) assay. With the two bispecific antibodies held constant at 10 nM and indinavir titrated, potent indinavir dose dependent T-cell activation was observed by CD69 upregulation and T-cell dependent cellular cytotoxicity (Fig.2c, 2d).

To demonstrate the ability of the IDV LITE Switch to control the assembly and activity of therapeutic proteins *in vivo*, we profiled the indinavir-dependent assembly of an active T-LITE in non-tumor-bearing C57BL/6 mice. The αCD3 and αEpCAM antibody fragments do not cross-react with their mouse homologs, so mice are a useful species to monitor pharmacokinetics without the confounding effects of target binding. Mice were dosed with 5, 1, or 0.2 mg/kg of each bispecific antibody intravenously in addition to 30 mg/kg indinavir sulfate also dosed intravenously. The total plasma concentration of the antibodies corresponded to the dose level and clearance was slow in the first 24 hours, as expected for Fc-containing antibodies (Fig.2f). In contrast, indinavir was cleared rapidly from the plasma after a single oral dose, consistent with previous reports (Fig.2g) ^15^. Mouse plasma samples were analyzed *ex vivo* in a co-culture assay that rapidly detects T-cell receptor activation by measuring phosphorylated SLP-76 (pSLP76). This assay provides a relative measurement of the amount of assembled T-LITE antibody complex present in the plasma samples. The degree of *ex vivo* T-cell activation tracked with the plasma concentration of indinavir (Fig.2h) despite the consistent levels of T-LITE antibodies over the 24-hour time period (Fig.2f), therefore demonstrating the reversible, indinavir-dependent properties of the system *in vivo*. In addition, formation of the antibody complex did not appear to qualitatively alter the clearance rate of the antibodies from expected clearance rates for human Fc-domain containing antibodies in mice. While binding to the LITE Switch complex did appear to extend the half-life of indinavir *in vivo*, we believe this would not meaningfully impact the reversibility of the system, as evidenced by the rapid decline of functional T-LITE complex in this experiment. Overall, these results support that the IDV LITE Switch has suitable properties to be utilized *in* vivo for reversible indinavir-dependent assembly and activation of a therapeutic.

In summary we present the engineering and characterization of a fully synthetic, bioorthogonal, humanized antibody-based CID system with optimal properties for therapeutic applications. We demonstrated through the application of a split T-cell engager that this tool can be utilized *in vitro* and *in vivo* to dose-dependently and reversibly regulate the assembly of proteins to cause a downstream biological signaling event. While this application highlights incorporation into recombinant therapeutic proteins to regulate signaling initiated by cell surface proteins, we believe the modular nature of this antibody-based CID system will facilitate broad adaptation into both intra- and extra-cellular systems, including cell and gene therapies. For intracellular applications, each of the switch domains can be readily reformatted to a single chain scFv format (sequences in Supplementary Materials) to function in the reducing environment of the cytoplasm while simultaneously minimizing the size of the genetic payload. In addition, the use of antibody domains should allow for N- or C-terminal fusion to proteins of interest while retaining binding, facilitating modularity in designing chemically regulated cellular signaling circuits. We envision the modular nature of the IDV LITE Switch will allow it to be used in a range of therapeutic applications including gene and cell therapies, as well as serving as a flexible research tool for the fields of cell, chemical, and synthetic biology.

## Acknowledgements

We would like to acknowledge Jim Wells, Peter Kim, Mike Brisken, Henry Lowman, Lawrence Fong, Art Weiss, and Brian Daniels for helpful scientific discussions.

## Author contributions

Experiments were conceptualized by: A.J.M., H.L.T., E.F.S., C.G., M.T., and Z.B.H. Experiments were performed by: H.L.T, A.L.C., Z.H., S.R., J.P.A., S.P., C.G., L.M., T.P., S.B., R.M., M.J., K.N., A.P., M.K., A.P., A.J.M., E.F.S. Data was analyzed by: A.J.M., H.L.T., E.F.S., A.L.C, S.P., C.G., L.M., S.B., R.M., K.N., M.T., Z.B.H. The manuscript was written by: A.J.M., H.L.T., J.P.A. The manuscript was edited by Z.B.H and E.F.S.

## Data availability statement

All data needed to evaluate the conclusions of the paper are present in the paper and/or the supplementary materials.

