## Supplemental Information for "A bioorthogonal antibody-based chemically-induced-dimerization switch for therapeutic application"

**Author Affiliations**

**Contents:**

I. **Methods**
II. **Supplementary Figures S1-S5**
III. **Flow Cytometry Gating Strategy**

### I. Methods

#### Vector Generation for Antibody Expression:

Plasmids for antibody expression were constructed by standard molecular biology methods. All DNA fragments were synthesized by IDT Technologies or Twist Biosciences and subcloned into either the pFUSE vector (InvivoGen) or pcDNA3.4 vector (Thermo Scientific) with Gibson assembly.

#### Antibody Expression and Purification:

All antibodies were expressed and purified from Expi293F (Thermo Fisher Scientific) or modified Expi293 BirA-KDEL (PMID: 29359686) cells according to an established protocol from the manufacturer (Thermo Fisher Scientific). Briefly, pFUSE (InvivoGen) vector encoding the protein of interest was transiently transfected into Expi293F cells at a fixed density of 3M/mL using the Expifectamine transfection kit and manufacturer's protocol (Thermo Fisher Scientific). Expression volumes varied, but a fixed vector:transfection reagent ratio (1mg : 2.8ml) was used. For multi-chain proteins, a stoichiometric equivalent of each respective vector was used. Culturing media for BirA cells was additionally supplemented with 100 µM of biotin prior to transfection for *in vivo* biotinylation. Enhancing supplements from the Expifectamine kit were added 20 hours post-transfection. Cells were incubated for a total of 4 d at 37 °C in an 8% $CO_2$ environment with orbital shaking before the supernatants were harvested by centrifugation. Fc-fusion proteins were purified by Protein A (MabSelect PrismA) affinity chromatography and His-tagged proteins were purified by Ni-NTA (Roche cOmplete™ His-Tag Resin) affinity chromatography. Eluted proteins from affinity purification were further separated by Size-exclusion chromatography with a Superdex 200 increase 10/300 GL column (Cytiva) in storage buffer (1X HBS + 5% Glycerol) as an aqueous phase. Samples were injected manually and fractionated via an AKTA Pure System with Software UNICORN 7.3 (Cytiva). Purity and integrity were assessed by SDS-PAGE with 4-12% BIS-TRIS precast gels (Thermo Fisher Scientific). Samples were stored at either 4°C or −80 °C with storage buffer in aliquots.

#### Small Molecule Reagents

Indinavir sulfate (>98%; Cayman Chemical #15150) was obtained commercially and used without further purification. Indinavir-NHS and Indinavir-Biotin were synthesized at WuXi AppTec and validated by NMR (Fig. S1).

#### Antibody discovery and humanization to generate LSA

Indinavir-NHS was conjugated to keyhole limpet hemocyanin (KLH) protein at a 40:1 Indinavir-NHS:KLH molar ratio and bovine serum albumin (BSA) protein at a 30:1 Indinavir-NHS:BSA molar ratio for 3 hours in phosphate buffer (100 mM phosphate, 150 mM NaCl, pH 8.0) followed by dialysis into PBS to generate Indinavir-KLH and Indinavir-BSA conjugates. Eight WT mice consisting of up to 3 strains were immunized with indinavir-KLH. By day 17 post-immunization, a high ELISA titer was observed against Indinavir-BSA compared to a negative control protein. A final boost consisting of a mixture of Indinavir-KLH and free Indinavir was given. The lymphocytes and splenocytes were collected and an immune scFv-phage library was constructed. The library underwent 3 rounds of panning against Indinavir-BSA conjugate. Top hits were reformatted as monovalent scFv-Fc fusions and screened in a BLI binding and competition assay to find antibodies where competition was observed in the presence of 100 nM soluble Indinavir (Fig. S2). Ab15 was humanized by grafting CDR sequences onto human VH1-46*01 and VK3-20*01 frameworks. LSA sequence is available in the Supplemental Materials (Fig. S3).

**Phage display selection to generate LSB**

Phage display selections were performed as previously described [1]. Briefly, selections with antibody phage Library E were performed using biotinylated-LSA captured with streptavidin-coated magnetic beads (Promega). Prior to each selection, the phage pool was incubated with 1 μM of biotinylated-LSA immobilized on streptavidin beads in the absence of Indinavir to deplete the library of any binders to the apo form of LSA. Subsequently, the beads were removed and Indinavir was added to the phage pool at a concentration of 10 μM. In total, four rounds of selection were performed with decreasing amounts of biotinylated-LSA antigen (100 nM, 50 nM, 25 nM, and 10 nM). LSB sequence is available in the Supplemental Materials (Fig.S3)

**Measurement of indinavir binding kinetics:**

The affinity of LSA to free Indinavir was measured by surface plasmon resonance (SPR) with a Biacore T200. The assay was performed at 25°C in 1x PBS-P+ buffer made from a 10x buffer stock (Cytiva 28995084). LSA in a scFv-Fc format was amine coupled to a Series S Sensor Chip CM5 (Cytiva 29149603). Trastuzumab, a non-binding control, was amine coupled to the reference lane. A single-cycle kinetics experiment was performed once with five sequential injections of 12.3, 37, 111, 333, 1000 nM Indinavir at 30 mL/min for 240 seconds each and dissociation was monitored for 600 seconds. The data was double referenced subtracted to the reference lane and five sequential injections of 0 nM Indinavir. Data was fit to a 1:1 Langmuir model to determine the association and dissociation rate constants.

**Measurement of LSA:LSB binding kinetics:**

The affinity of LSB binding to LSA was measured by SPR with a Biacore T200. The assay was performed at 25°C in 1x HBS-EP+ buffer made from a 10x buffer stock (Cytiva BR100669). A monovalent LSA scFv-Fc fusion was captured on a Series S Sensor Chip CM5 (Cytiva 29149603) using the Human Antibody Capture Kit (Cytiva BR100839). For Indinavir dependent binding, a multi-cycle kinetics experiment was run with 0 nM, 3.7, 11.1, 33.3, 100, and 300 nM LSB Fab in the presence of 10 mM Indinavir for 300 seconds at 30 mL/min followed by a dissociation step for 600 seconds. For kinetics without Indinavir, a similar multi-cycle kinetics experiment was run at 0, 100, 300, and 900 nM without Indinavir present. Each binding experiment was performed once. The data was double referenced subtracted to the reference lane and the 0 nM sensogram. The sensor chip was regenerated with an injection of 3 M magnesium chloride for 45 seconds at 30 mL/min after each cycle. Data was fit to a 1:1 Langmuir model to determine the association and dissociation rate constants.

**BLI Indinavir Washout Experiment**
LITE switch Indinivir washout/reversibility kinetics were measured by biolayer interferometry (BLI) on an Octet® Qke (Sartorius) instrument. Biotinylated-LSA scFv-Fc, in the presence of 1 μM Indinavir or 0.05% DMSO vehicle, was immobilized on streptavidin biosensors and loaded to a signal of 1.2-1.8 nm. After loading, biosensors were blocked with 10 μM biotin containing 1 μM Indinavir or vehicle. Association steps were performed for 10 minutes in the presence of 50 nM LSB Fab and 1 μM Indinavir or vehicle. At steady-state binding of LSB to LSA, LITE switch complexes dissociated for 2 hours in buffer containing: (1) [No washout] 50 nM LSB and 1 μM IDV, (2) [Indinavir washout] 50 nM LSB (3) [Indinavir and LSB washout] buffer only. Each experiment was performed once.

**Complex Formation by Size Exclusion Chromatography**
To assess complex formation in solution, a size exclusion chromatography (SEC) assay was performed on a Superdex 200 increase 10/300 in PBS pH 7.4 buffer. In three separate runs, 100 mg LSA + 5 mM Indinavir, 100 mg LSB, and 100 mg LSA + 100 mg LSB + 5 mM Indinavir were injected onto the SEC column and the absorbance was monitored at 280 nm. Data from a single representative run is reported. The earlier retention time of the LSA:IDV:LSB sample demonstrates a high degree of complex formation. For these experiments the following mutational variants of LSA and LSB were utilized: LSA H40A M50I N59S, LSB A50S Y102W A109L. These residues were later mutated as part of optimization of affinity and stability.

**Negative Stain Electron Microscopy**
Negative stain EM experiments were performed on a Tecnai T12 120 kV microscope with a 1.68 Å/pixel size. 2% uranyl acetate was used as the stain. A formvar carbon supported copper grid with a mesh size of 400 was used for molecular adsorption. The LSA:IDV:LSB complex was purified by SEC, adsorbed onto the grid, underwent wash and stain steps, then proceeded to image collection and data analysis. A representative micrograph is shown in Fig. S4a. 2D class averages were generated and shown in Fig. S4b. 50K particles were used to generate a 3D EM map of the LSA:IDV:LSB complex. Homology models of LSA and LSB Fabs were generated in Schrödinger BioLuminate software and fit into the 3D EM density showing the CDR regions coming into contact. For this experiment the following mutational variants of LSA and LSB were utilized: LSA H40A M50I N59S, LSB A50S Y102W A109L.

**AlphaLISA assay**
AlphaLISA bead mixtures were prepared in HiBlock buffer by combining anti-FLAG Acceptor Beads (40 μg/mL), Biotinylated CD3-Fc (10 nM) and FLAG-EpCAM (10 nM) in a volume of 10 uL per well of a ProxiPlate-384 Plus, White 384-shallow well Microplate (PerkinElmer). To this, 5 μL of Streptavidin Donor Beads (80 μg/mL) were added, and the bead mixture was incubated overnight at 4℃. The following morning, the mouse plasma samples (5 μL) were added to the bead mixture and incubated at room temperature for 90 minutes. Plates were read on a Perkin-Elmer EnVision plate reader with Alpha module

(615 nm detection). The data reported in Fig.2a shows the mean from a single experiment run in triplicate with error bars representing standard deviation.

**Isolation of T-cells for in vitro assays**

Leukapheresis packs were obtained from deidentified healthy adult donors (StemCell Technologies, Vancouver, BC). Peripheral blood mononuclear cells (PBMCs) were first isolated using the EasySep Direct Human PBMC Kit (StemCell Technologies). Human T cells were magnetically isolated from the PBMCs using the EasySep Human T cell Isolation Kit (StemCell Technologies). T cells were cryopreserved in DMEM with 10% FBS and 10% DMSO and thawed the day of experiments.

**T-cell activation and T-cell dependent cellular cytotoxicity assays**

Co-culture assays were used to assess T-cell activation (CD69 upregulation), cytokine production or T-cell-dependent cellular cytotoxicity (TDCC) of target cells. Flat-bottom 96-well plates were seeded with 5,000 HCT-116 human colon carcinoma (ATCC Cat# CCL-247) target cells in complete growth media (DMEM + 10% FBS + 1% penicillin/streptomycin) 16-24 hrs before the assay. Antibody drugs were added in 10 µL growth media and incubated for at least 10 min at 37°C. Indinavir or vehicle control was added in 10 µL growth media. Magnetically isolated human T-cells were thawed, washed, and resuspended in complete growth media, then added to target cells (50,000 T cells per well) in 80 µL growth media for a final assay well volume of 100 µL. The final effector:target (E:T) ratio was 10:1. Assay plates were incubated at 37°C with 5% $CO_2$ for 66 - 70 hr depending on the experiment. T-cells were transferred to a U-bottom 384-well plate and spun (600 rcf x 5 min). Supernatants were stored at -20°C for later analysis of cytokines. T-cells were resuspended in CF405M viability dye (0.5 µM, Biotium, Fremont, CA) in PBS for 10 min at 37°C. At the end of viability staining, cells were fixed by adding methanol-free paraformaldehyde (Electron Microscopy Sciences, Hatfield, PA) to a final concentration of 1% for 10 min at room temperature. T-cells were spun (600 rcf x 5 min) and supernatant was aspirated. T-cells were stained with FITC-conjugated anti-human CD45 (BioLegend, San Diego, CA) at 1 µg/mL final dilution and PE-conjugated anti-human CD69 (BioLegend, San Diego, CA) at 900 ng/mL final dilution for 30 minutes at room temperature in the dark, then immediately run an IntelliCyt® iQue3 flow cytometer (Sartorius AG, Göttingen, Germany). While T cells were being processed, target cells were incubated washed with PBS and resuspended in 50 µL growth media then mixed with 20 µL CellTiter-Glo 2.0 (Promega, Madison, WI) for 15 min at room temperature with shaking then read for luciferase light emission on a GloMax Explorer plate reader using default settings. Specific Cytotoxicity (%) was calculated from the relative luminescence unit values (RLU) using the formula: [ 1 – ( experimental RLU / vehicle-treated RLU ) ) x 100 ]. The frequency of CD69+ T-cells (%) was calculated as the percentage of CD69-positive events within the CF405M-negative population. Dose-response curves and $EC_{50}$ values were calculated in GraphPad Prism 9 software (GraphPad Software San Diego, CA). The data reported in Fig.2c and Fig.2d shows the mean from a single experiment run in triplicate with error bars representing standard error of the mean.

**In vivo PK and T-LITE activation**

Pharmacokinetics experiments were performed under an approved IACUC protocol (Protocol # CR-0178). Female C57BL/6 mice, aged 7-8 weeks, were purchased from Charles River. Antibody test articles were formulated on the morning of injection. For T-LITE antibodies, TAAxLSA and CD3xLSB antibodies were stored as separate stocks until the moment at which they were co-formulated. Formulations were diluted with phosphate buffered saline without calcium or magnesium, pH 7.4 (Corning Cat# 21-040) to the required concentrations, then stored on ice. Indinavir sulfate was formulated up to 7 days prior to injection by mixing indinavir sulfate (Cayman Chemical Cat# 15150) with Non-lactated Standard Ringer Solution (BioWorld Cat# 40120236-1) at 6.96 mg/mL (6 mg/mL active drug). Mice (n=3 per time point per group) were injected at 5 mL/kg in the lateral tail vein using an insulin syringe (BD Cat# 329420). Injections were staggered to ensure accurate timing between injections and blood collection. Blood was collected by cardiac puncture under deep isoflurane anesthesia into BD Microtainer K2 EDTA tubes (BD Cat# 365974) using a 25G syringe (BD Cat# 309626). Blood was centrifuged for 10 minutes at 2000 x g to isolate plasma which was stored frozen at -80°C.

**Anti-human IgG1 quantitation by ELISA**

A universal anti-human IgG1 ELISA was used to measure total T-LITE antibody concentration in mouse plasma. A goat anti-human IgG capture antibody (Southern Biotech #2049-01) and a mouse anti-human IgG Fc-HRP detection antibody (Southern Biotech #9040-05, Clone JDC-10) were used in this assay. ELISA reagents were sourced from Biolegend: ELISA Coating Buffer (5X) (#421701), ELISA Assay Diluent (5X) (#421203), ELISA Wash Buffer (20X) (#421601), ELISA Substrate Solution A and B (#421101), and ELISA Stop Solution (#423001). Capture antibody was diluted to 5 ug/mL in 1X coating buffer, and 100 µl was added to a 96-well flat bottom MaxiSorp Immuno Plate (Nunc #442404). The capture antibody was incubated for one hour at 37 °C in a humidified incubator and then washed three times with 200 µl of 1X wash buffer. 100 µl of 1X assay diluent was added to each well and incubated for one hour at room temperature, and then the plate was washed three times with 200 µl of 1X wash buffer. Standard curves composed of equimolar concentrations of T-LITEs A and B in mouse plasma (Rockland #D508-05-0100) were made with total antibody concentrations between 0.25 – 16 nM. Samples with high concentrations of antibody outside of the range of the standard curve were diluted in mouse plasma. 60 µl of standards and plasma samples were diluted 1:4 in 1X assay diluent, and 100 µl were added to the plate in duplicate. Samples were incubated in the plates for two hours at room temperature and then washed five times with 1X wash buffer. The detection antibody was diluted 1:32000 in 1X assay diluent, and 100 µl of diluted detection antibody was added to each well of the plate. The plate was incubated for one hour at room temperature and then washed seven times with 200 µl 1X wash buffer, allowing the buffer to soak on the plate for 30 seconds/wash. The TMB substrate working solution was made with equal volumes of Substrate Solutions A and B, and 100 µl of TMB substrate working solution was added to each well. The plate was incubated for 30 minutes at room temperature in the dark. 50 µl of stop solution was added, and the absorbance read at 450 nm wavelength. A comparability study comparing standard curves of T-LITE A and T-LITE B by themselves provided signals within ~20% of the equimolar T-LITE A and B standard curve, indicating that this standard curve could be used even for samples with an overabundance of one T-LITE antibody over the other.

**LC-MS quantitation in mouse plasma**

An LC-MS based PK assay was developed and qualified for measuring the total amount of indinavir in plasma samples. In this assay, 25 µl of calibration standards (0.6-600 ng/mL indinavir, Cayman Chemicals #15150, in cyno or mouse plasma sourced from BioIVT), QC's, plasma blanks, or plasma samples were added to a 96-well deep well plate. Samples with high concentrations of indinavir outside of the range of the standard curve were diluted in plasma. Next, 25 µl of internal standard (IS) working solution (50 ng/mL indinavir-d6, Cayman Chemicals #29585, in 50:50 water:acetonitrile) and 400 µl water were added to each well. The plate was vortexed at 1650 rpm for 3 mins. 200 µl from each well was transferred to a Biotage ISOLUTE SLE+ 400 µl 96-well plate and allowed to adsorb to the plate for 10 mins. Indinavir was eluted from the plate with two aliquots of 500 µl methyl t-butyl ether (MTBE), and the eluate was evaporated to dryness under a stream of nitrogen gas at a temperature of 50 °C. The extracted indinavir was reconstituted with 200 µl of a solution containing 10% acetonitrile and 90% 0.02% formic acid and 10 mM ammonium formate in water. The reconstituted indinavir samples were vortexed at 1800 rpm for one minute. 50 µl of these samples were added to another 96-well plate containing 200 µl of solution containing 10% acetonitrile and 90% 0.02% formic acid and 10 mM ammonium formate in water, and the samples were again vortexed at 1800 rpm for one minute before being placed in the LC-MS autosampler.

Samples were analyzed on a Sciex API 6500+ triple quadrupole mass spectrometer equipped with a Shimadzu Prominence UHPLC. 2 µl of each sample was injected onto a Waters Acquity BEH C8 2.1 x 50 mm, 1.7 um column at a flow rate of 0.6 mL/min and a gradient of 20% B to 95% B over 1.10 minutes. Mobile Phase A was 0.02% formic acid and 10 mM ammonium formate in water (pH 4.1), and Mobile Phase B was 0.1% formic acid in acetonitrile. Indinavir transition 614.4 m/z à 421.4 m/z and indinavir-d6 (IS) transition 620.5 m/z à 421.4 m/z were used to quantify the levels of indinavir in each sample.

The assay was qualified over a standard curve range of 0.6 – 600 ng/mL and QC levels of 0.6 ng/mL (LLOQ), 1.8 ng/mL (LQC), 75 ng/mL (MQC), and 450 ng/mL (HQC). %CV and %Accuracy for all QC levels were within 20%. %CV and %Accuracy for QC standards were also within 20% after a three-day autosampler stability study at room temperature. There was no background signal in blank plasma samples, and LLOQ measurement %Accuracy was within 20% in six different lots of cyno plasma.

For metabolite analysis, the same protocol was used, but indinavir and three metabolites were quantified in the same assay. Three metabolite standards, M3, M5, and M6, as well as d5-labelled versions of these three metabolites were synthesized at WuXi. Equal concentrations of indinavir and metabolites were added to each calibration or QC standard, and equal concentrations of indinavir-6 and d5-labelled metabolites were added to the IS working solution. The transitions in Table ## were used to quantify indinavir and metabolites in each sample. The assay was qualified over a standard curve range of 0.6 – 600 ng/mL and QC levels of 0.6 ng/mL (LLOQ), 1.8 ng/mL (LQC), 100 ng/mL (MQC), and 450 ng/mL (HQC). %CV for all QC levels were within 20%, except for M6 LLOQ, which had a %CV within 40%. %Accuracy for all QC levels were within 35% except for M6 LLOQ, which showed high variability.

**pSLP76 bioassay**
A co-culture bioassay measuring T-cell activation via a phospho-SLP-76 AlphaLISA assay was used to detect T-LITE complex in plasma. Standard curves were generated by titrating 10x T-LITE in mouse serum containing 10x small molecule (Indinavir). This was incubated for 30 minutes at 37 °C in a humidified incubator to form the T-LITE complex. Jurkat cells (ATCC #TIB-152, Clone E6.1) and HCT-116 cells (ATCC #CCL-247) were grown according to ATCC recommendations and combined in a 5:1 ratio (44.4e6 Jurkat cells:8.8.9e6HCT-116 cells in 20 mL) in DMEM (Corning #10-013-CM) without serum. 90 µl of cell mixture was added to each well in 96-well U-bottom plates (ThermoFisher #168136) and pelleted at 400 xg for one minutes at room temperature. Next, 10 µl of each plasma sample, the pre-formed T-LITE complex or a positive control lysate were added to the plate in singlicate, and the plate was incubated for 15 minutes at 37 °C in a humidified incubator. Cells were pelleted again at 400 xg for five minutes at room temperature and media was aspirated. The remainder of the protocol used an AlphaLISA Surefire Ultra pSLP76 (Ser 376) kit (PerkinElmer #ALSUP-PSLP-A10K). 75 µl of 1X lysis buffer was added to each well and shaken at 350 rpm for 20 minutes at room temperature. 30 µl of lysate per well was transferred to a half-area Optiplate (PerkinElmer #6002290). 15 µl of Acceptor bead mix was added to each well. The plate was shaken at 350 rpm for two minutes at room temperature and then incubated for one hour at room temperature in the dark. 15 µl of Donor bead mix was added to each well. The plate was vortexed at 350 rpm for two minutes at room temperature and then incubated for two hours at room temperature in the dark. The plate was read on an EnVision microplate reader (PerkinElmer)using standard AlphaLISA settings.

1. Hill, Z. B., Martinko, A. J., Nguyen, D. P. & Wells, J. A. Human antibody-based chemically induced dimerizers for cell therapeutic applications. *Nat Chem Biol* 14, 112–117 (2018).

### <u>II. Supplementary Figures S1-S5</u>

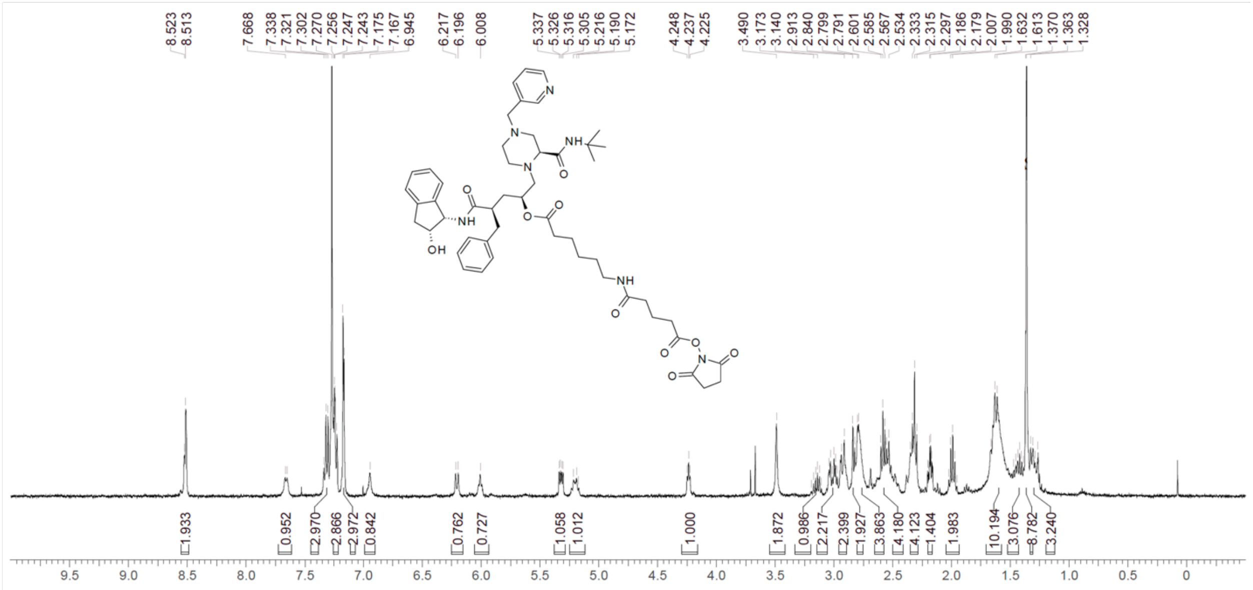

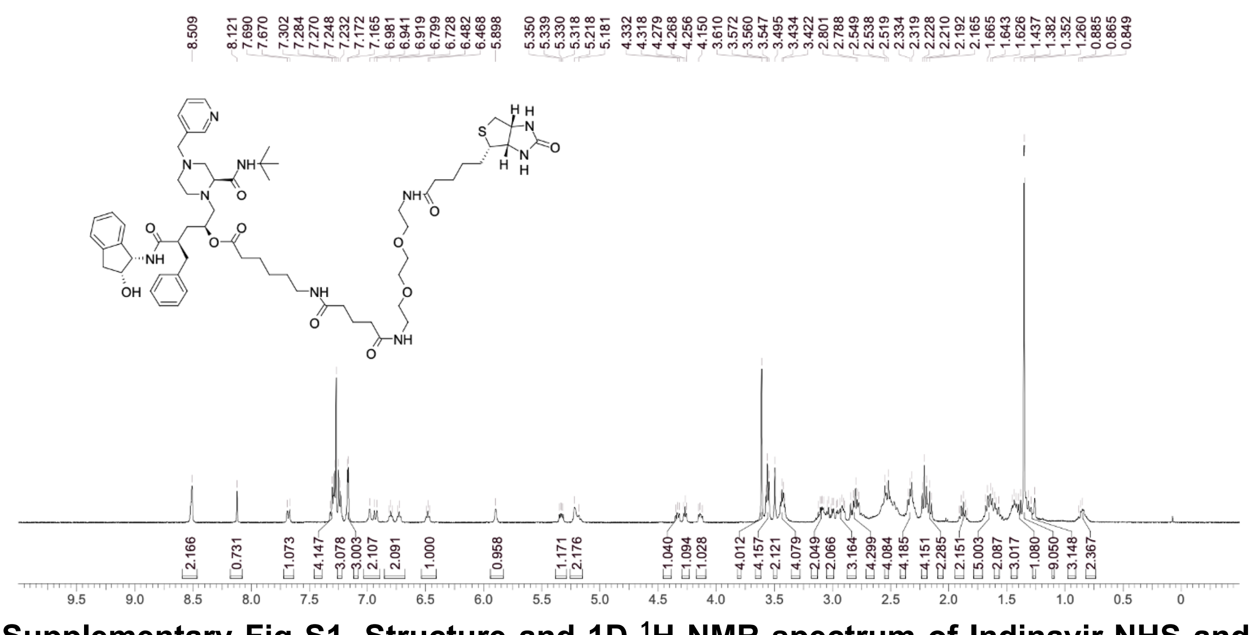

**Supplementary Fig S1. Structure and 1D $^{1}$H NMR spectrum of Indinavir-NHS and Indinavir-Biotin** Indinavir-NHS (Top) and Indinavir-Biotin (Bottom) were synthesized at WuXi AppTec, dissolved in CDCl3, and a 1D $^{1}$H NMR spectrum was taken on a 400 MHz Bruker NMR to validate the structure.

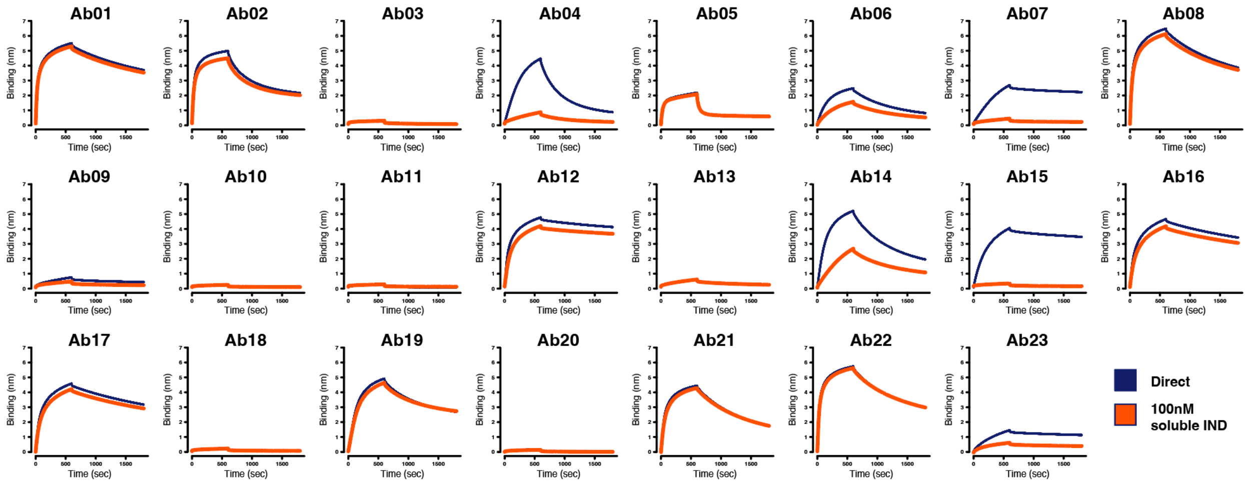


**Supplementary Fig S2. Indinavir antibody binders profiled by BLI after phage display selection**. 23 antibodies were profiled for direct indinavir binding and competition. Ab15 showed competition with soluble indinavir and was selected for humanization resulting in the LSA antibody fragment.

- LSA Fab
  - Heavy Chain:
    QVQLVQSGAEVKKPGASVKVSCKASGYTFTSYWMHWVRQHPGQGLEWMGMIHPNSGGTNYAQKFQGRVTMTRDTSTSTVYMELSSLRSEDTAVYYCAMGDYVSRDY
    WGQGTLVTVSSASTKGPSVFPLAPSSKSTSGGTAALGCLVKDYFPEPVTVSWNSGALTSGVHTFPAVLQSSGLYSLSSVVTVPSSSLGTQTYICNVNHKPSNTKVDKKVE
    PKSC
  - Light Chain:
    EIVLTQSPGTLSLSPGERATLSCRASSSVSSSYLHWYQQKPGQAPRLLIYSTSSRATGIPDRFSGSGSGTDFTLTISRLEPEDFAVYYCQQYSGYQFTFGQGTKLEIKRTVA
    APSVFIFPPSDEQLKSGTASVVCLLNNFYPREAKVQWKVDNALQSGNSQESVTEQDSKDSTYSLSSTLTLSKADYEKHKVYACEVTHQGLSSPVTKSFNRGEC
- LSA scFv
  - QVQLVQSGAEVKKPGASVKVSCKASGYTFTSYWMHWVRQHPGQGLEWMGMIHPNSGGTNYAQKFQGRVTMTRDTSTSTVYMELSSLRSEDTAVYYCAMGDYVSRDY
    WGQGTLVTVSSGGGGSGGGGSGGGGSEIVLTQSPGTLSLSPGERATLSCRASSSVSSSYLHWYQQKPGQAPRLLIYSTSSRATGIPDRFSGSGSGTDFTLTISRLEPEDF
    AVYYCQQYSGYQFTFGQGTKLEIK

- LSB Fab
  - Heavy Chain:
    EVQLVESGGGLVQPGGSLRLSCAASGFDFSSYSIHWVRQAPGKGLEWVAAISPYYGSTYYADSVKGRFTISADTSKNTAYLQMNSLRAEDTAVYYCARYYEYYKYDLYAG
    SLGFDYWGQGTLVTVSSASTKGPSVFPLAPSSKSTSGGTAALGCLVKDYFPEPVTVSWNSGALTSGVHTFPAVLQSSGLYSLSSVVTVPSSSLGTQTYICNVNHKPSNTK
    VDKKVEPKSC
  - Light Chain:
    DIQMTQSPSSLSASVGDRVTITCRASQSVSSAVAWYQQKPGKAPKLLIYSASSLYSGVPSRFSGSRSGTDFTLTISSLQPEDFATYYCQQGGYSLITFGQGTKVEIKRTVAA
    PSVFIFPPSDSQLKSGTASVVCLLNNFYPREAKVQWKVDNALQSGNSQESVTEQDSKDSTYSLSSTLTLSKADYEKHKVYACEVTHQGLSSPVTKSFNRGEC

- LSB scFv
  - EVQLVESGGGLVQPGGSLRLSCAASGFDFSSYSIHWVRQAPGKGLEWVAAISPYYGSTYYADSVKGRFTISADTSKNTAYLQMNSLRAEDTAVYYCARYYEYYKYDLYAG
    SLGFDYWGQGTLVTVSSGGGGSGGGGSGGGGSDIQMTQSPSSLSASVGDRVTITCRASQSVSSAVAWYQQKPGKAPKLLIYSASSLYSGVPSRFSGSRSGTDFTLTISS
    LQPEDFATYYCQQGGYSLITFGQGTKVEIK

**Supplementary Fig S3**. **Sequences of LSA and LSB Indinavir-LITE switch antibodies.** Sequences of LSA and LSB antibody fragments used in this study. Sequences are provided as Fab and scFv format and can be incorporated into antibodies, protein fusions, or genetic fusions.

**A** Representative micrograph
Fab1-Fab2-SM complex (c = 0.003 μg/μL)

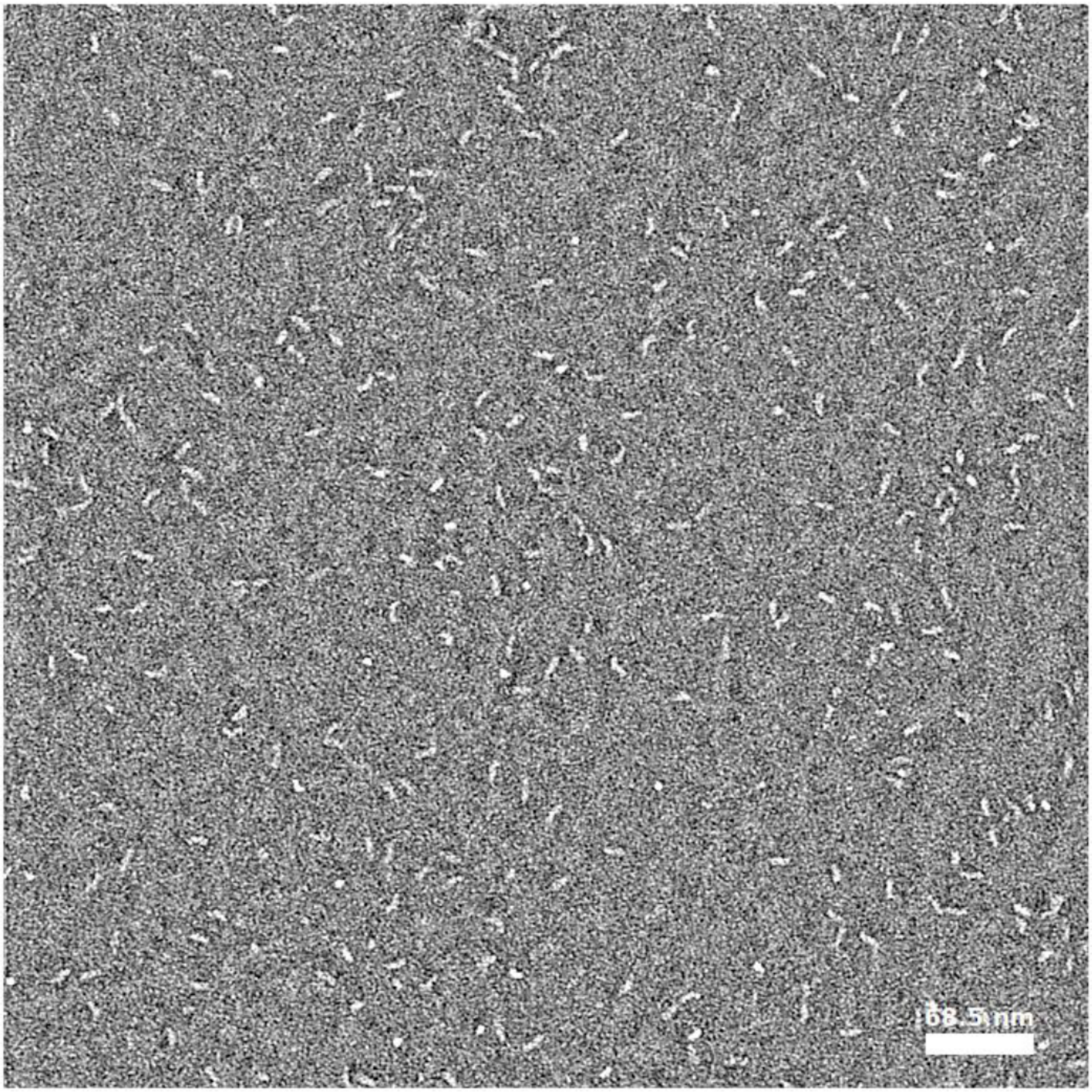


No aggregation of particles

Scale bar = 68.2 nm

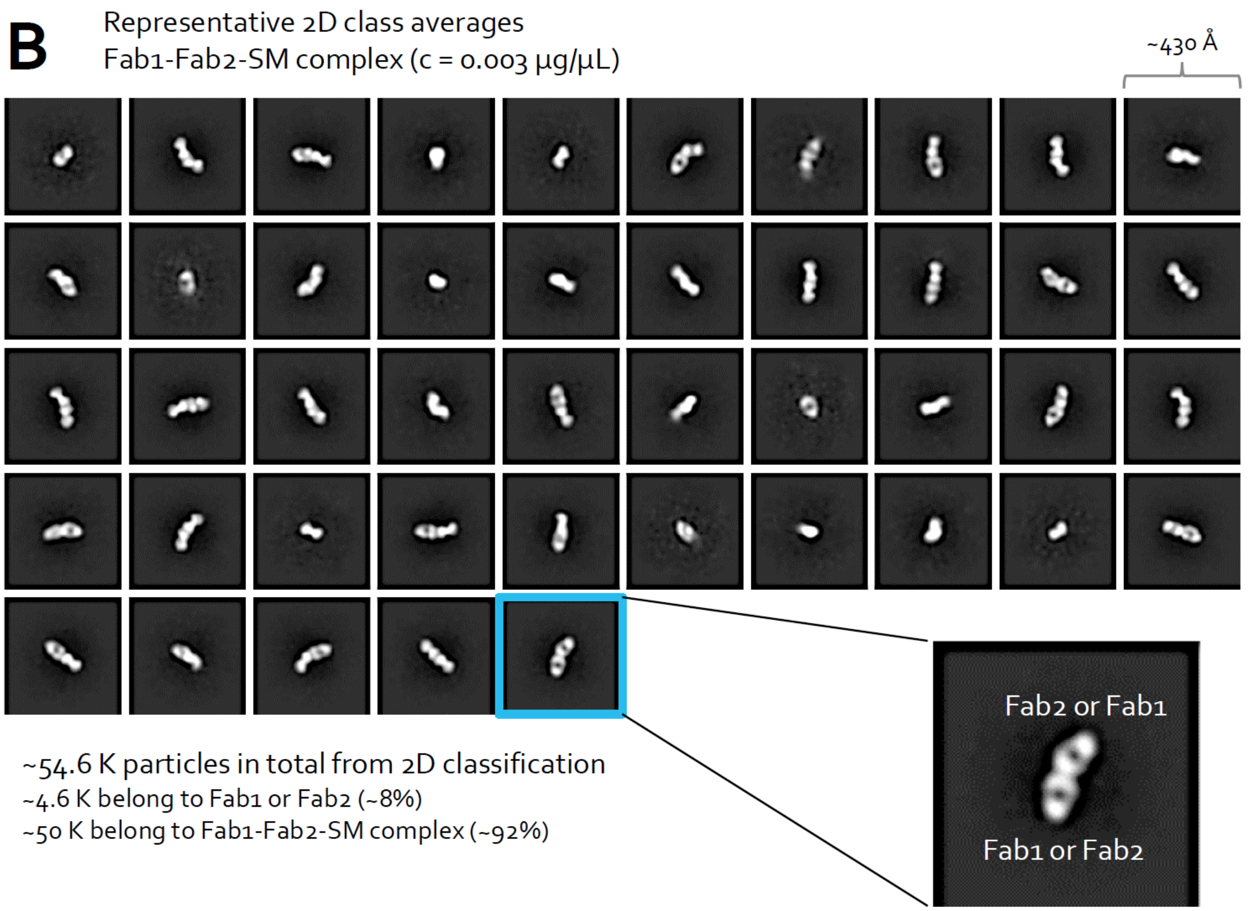


**Supplementary Fig S4. Negative stain electron microscopy shows the formation of the T-LITE switch.** (A) A representative micrograph of the LSA-LSB-IDV complex (B) Representative 2D class averages of the LSA-LSB-IDV complex in various orientations used to assemble a 3D map.

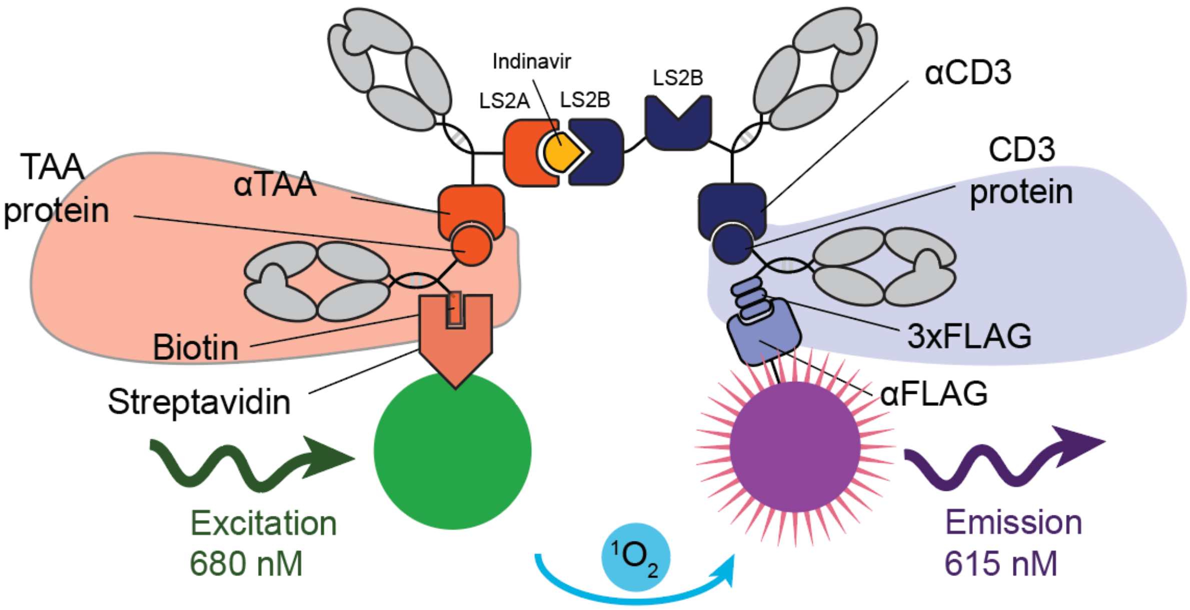


**Supplementary Fig S5. AlphaLISA assay to detect CD3-EpCAM interaction.** Biotin (L) and CD3-FLAG (M) detection molecules designed to bind to the bispecific antibody (A3,B3). AlphaLISA beads were directed against the biotin and Flag tags on the detection molecules. Excitation of the donor with a light source induces production of singlet oxygen that diffuses toward the acceptor beads and starts an electron transport process that is converted to light energy by acceptor chemistry. It is read at 615 nm and is characterized as an Alpha signal. Therefore, the specific Alpha signal at 615 nm is proportional to the amount of CD3/αTAA bispecific antibody present in the sample. The assay format used is a sandwich assay, as shown above.

### III. Flow Cytometry Gating Strategy

The gating strategy to identify CD69-positive viable T-cells in Fig.2c was as follows: (1) "Cells" were defined as FSC-H(mid) SSC-H(mid) with the gate drawn to exclude bead controls that were spiked into the sample. (2) Within "Cells", "Singlets" were defined based on correlation of FSC-A and FSC-H, with the gate drawn to exclude FSC-H(high) doublets. (3) Within "Singlets", CF405M-high dead cells were excluded by gating on CF405M(low) to define "Viable Tcells" events. (4). Within "Viable Tcells", the cells with upregulated CD69 were gated as CD69(high) to define "CD69+ Viable T-cells". The abundance of "CD69+ Viable T-cells" was quantified in each sample and graphed as a bar plot.

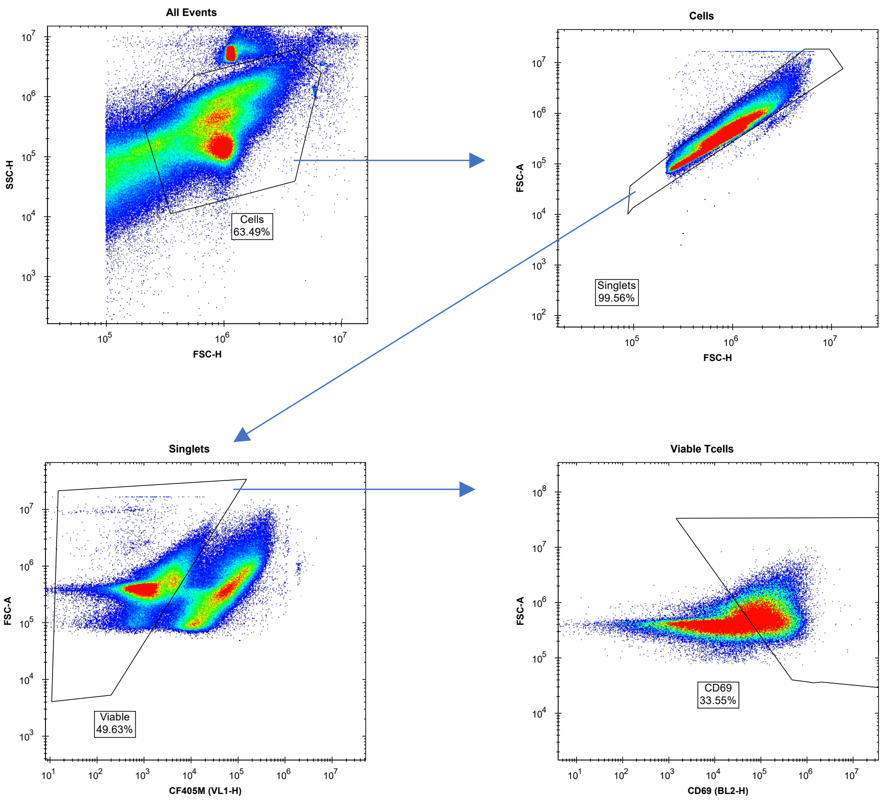
